# Symmetrically-dispersed spectroscopic single-molecule localization microscopy

**DOI:** 10.1101/2019.12.14.876557

**Authors:** K. Song, Y. Zhang, B. Brenner, C. Sun, H. F. Zhang

## Abstract

Spectroscopic single-molecule localization microscopy (sSMLM) achieved simultaneously imaging and spectral analysis of single molecules for the first time. Current sSMLM fundamentally suffers from reduced photon budget because of dividing photons from individual stochastic emission into spatial and spectral channels. Therefore, both spatial localization and spectral analysis only use a portion of the total photons, leading to reduced precisions in both channels. To improve the spatial and spectral precisions, we present symmetrically-dispersed sSMLM or SDsSMLM to fully utilize all photons from individual stochastic emissions in both spatial and spectral channels. SDsSMLM achieved 10-nm spatial and 0.8-nm spectral precisions at a total photon budget of 1000. Comparing with existing sSMLM using a 1:3 splitting ratio between spatial and spectral channels, SDsSMLM improved the spatial and spectral precisions by 42% and 10%, respectively, under the same photon budget. We also demonstrated multi-color imaging in fixed cells and three-dimensional single-particle tracking using SDsSMLM.

## Introduction

The ability of spectroscopic single-molecule localization microscopy (sSMLM) to capture spectroscopic signatures of individual molecules together with their spatial distribution allows observing subcellular structure and dynamics at nanoscale. As a result, sSMLM has shown great potential in understanding fundamental biomolecular processes in cell biology and material science (*1*–*10*). It also enables characterizing nanoparticle properties based on the emission spectrum at the single-particle level (*11*–*15*). Similar to other localization-based super-resolution techniques, such as stochastic optical reconstruction microscopy (STORM), photoactivated localization microscopy (PALM), and point accumulation for imaging in nanoscale topography (PAINT), the localization precision of sSMLM is fundamentally limited by the number of collected photons per emitter (*16*–*18*). However, sSMLM suffers from further photon budget constraint since the collected photons from each stochastic emission need to be divided into two separate channels to capture the spatial and spectral information simultaneously (*7*–*14*). Thus, spatial localization precision of sSMLM also depends on the splitting ratio between the spatial and spectral channels, and is typically limited to 15-30 nm in cell imaging (*3*–*8*). Although a dual-objective sSMLM design was previously demonstrated with improved spatial localization precision, it imposes a constraint on live cell imaging and adds complexity to system alignment (*1*, *19*). Splitting photons into two channels in sSMLM forces an inherent trade-off between spatial and spectral localization precisions (*7*). Currently, there lacks a method to fully utilize the full photon budget to maximize both spatial and spectral localization precisions in sSMLM.

To overcome this inherent trade-off, we develop the symmetrically-dispersed sSMLM or SDsSMLM, which has two symmetrically-dispersed spectral channels instead of one spatial and one spectral channels. SDsSMLM fully utilizes all collected photons for both spatial localization and spectral analysis. We showed improvements in spatial and spectral localization precisions via numerical simulation and validated them through imaging fluorescent nanospheres and quantum dots (QDs). We further demonstrated multi-color imaging of subcellular structures and three-dimensional (3D) single-particle tracking (SPT) capabilities.

## Results

### SDsSMLM

The concept of SDsSMLM is illustrated in Fig. 1. SDsSMLM is based on a conventional single-molecule localization microscopy (SMLM) system with an added grating-based spectrometer (Fig. S1). In the emission path, the fluorescence light is confined by a slit at the intermediate image plane and symmetrically dispersed into the −1^st^ and 1^st^ orders by a transmission grating with equal splitting ratio (Fig. 1A). Then, these dispersed fluorescence emissions are captured by an electron multiplying charge-coupled device (EMCCD) camera to form two symmetrical spectral images after passing through the relay optics.

**Fig. 1.**
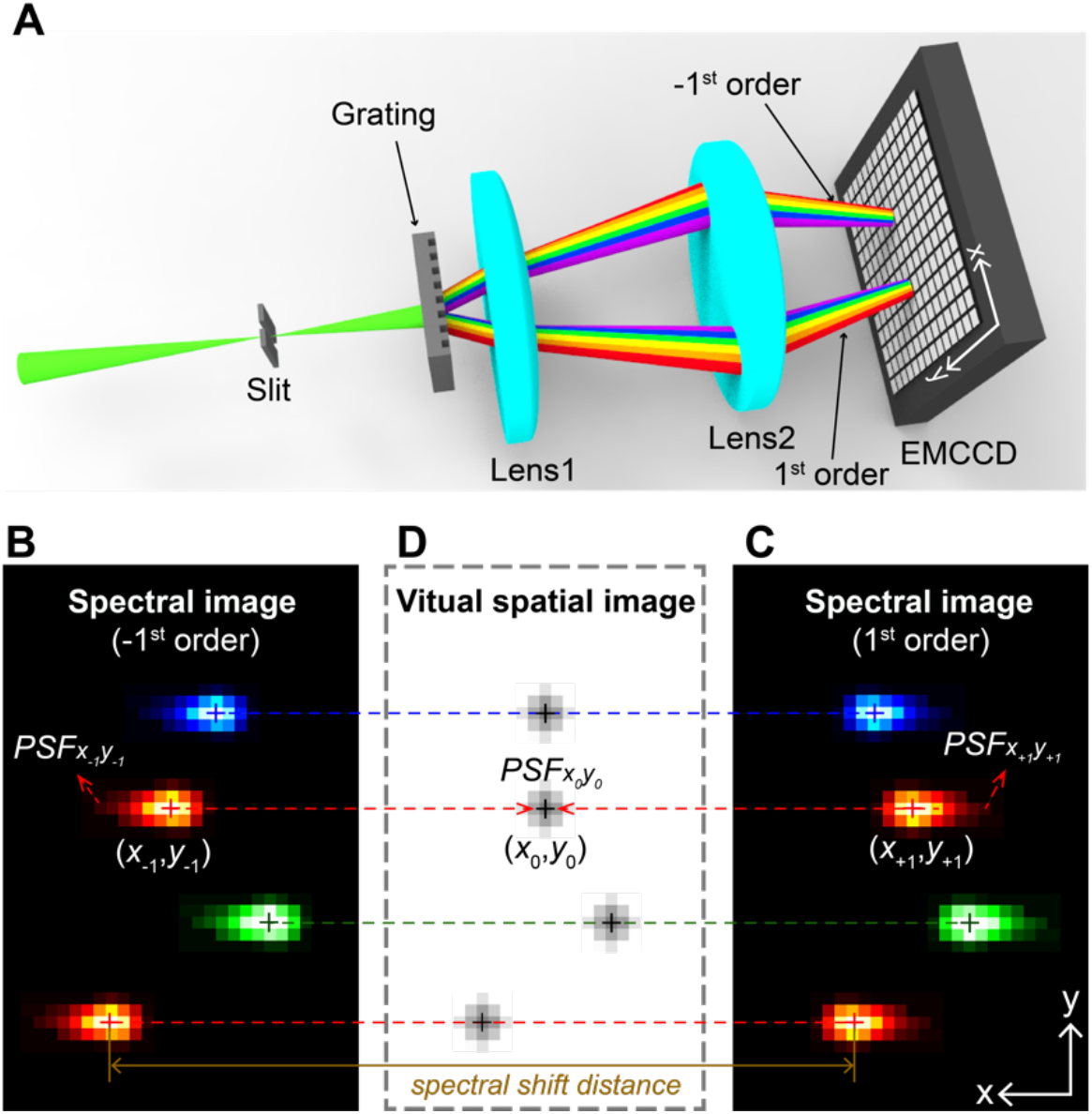
Working principle of SDsSMLM. **(A)** Schematic of SDsSMLM experimental system; **(B)** Illustrative image containing four single molecules from the −1^st^ order spectral channel; **(C)** Corresponding image of the same four molecules from the 1^st^ order spectral channel; **(D)** Calculated virtual spatial image from the two spectral images shown in panels B and C.

While existing sSMLM simultaneously captures spatial (0^th^ order) and spectral (1^st^ order) images, SDsSMLM captures only two spectral images (−1^st^ and 1^st^ orders, Figs. 1B & 1C) of individual molecules. The two spectral images of a particular single molecule emission are mirror images of each other with respect to the true location of the molecule. Therefore, we can localize single molecules by identifying the middle points (black plus symbols in Fig. 1D) between the two symmetrically-dispersed spectral images. This symmetry-middle point relationship holds true for all molecules regardless of their emission spectra and minute spectral variations even among the same species of molecules. Identifying all the middle points will generate a virtual spatial image (Fig. 1D). This virtual spatial image utilizes all the detected photons in each EMCCD frame as compared with a portion of the photons in existing sSMLM. It also should be noted that the virtual spatial image is not affected by the spectral heterogeneity of individual molecules, which is cancelled out through the symmetry-middle point relationship.

In SMLM, we estimate the localization position of individual molecules in the spatial image with a limited certainty (*20*). When the localization position is estimated repeatedly, the spatial localization precision (referred to as the spatial precision) is described as the standard deviation of the distribution of the estimated localization positions. Similarly, in SDsSMLM, we estimate the localization positions (*x*_−1_, *y*_−1_) and (*x*_+1_, *y*_+1_) from the −1^st^ order and 1^st^ order spectral images (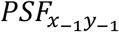 and 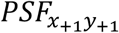 in Figs. 1B & 1C). Then, we determine the localization position (*x*_0_, *y*_0_) in the virtual spatial image (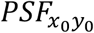 in Fig. 1D) using (*x*_−1_, *y*_−1_) and (*x*_+1_, *y*_+1_), as shown in Figs. 1B-D. Accordingly, the spatial precision in SDsSMLM is described by the standard deviation of the distribution of the estimated (*x*_0_, *y*_0_) in the virtual spatial image (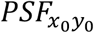).

In addition, from the two spectral images (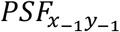 and 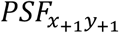), we generate new spectral images (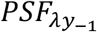 and 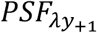) based on spectral calibration (details are described in Methods and Materials.). Then, we integrate them along the y-axis and extract spectral centroids (*λ*_*SC*_) to represent emission spectra of individual molecules. We calculate *λ*_*SC*_ as *λ*_*SC*_ = ∑_*λ*_ *λI*(*λ*) / ∑_*λ*_ *I*(*λ*), where *λ* is the emission wavelength and *I*(*λ*) is the spectral intensity at *λ* (*7,23*). Accordingly, the spectral localization precision (referred to as the spectral precision) is described as the standard deviation of the spectral centroid distribution.

Specifically, to generate the virtual image, we first localize the two spectral images (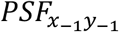 and 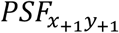) of individual molecules along the x-axis using Gaussian fitting based on maximum likelihood estimators (MLE), which achieves the precision closest to the Cramér-Rao Lower Bound (*17*, *18*, *21*, *22*). Then, we obtain the two localization positions *x*_−1_ and *x*_+1_, which are symmetrically distributed with respect to the true location of the molecule. Therefore, we can determine the spatial location *x*_0_ in the virtual image by calculating the mean value of *x*_−1_ and *x*_+1_. In addition, we localize the two spectral images (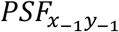 and 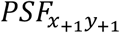) along the y-axis, which generates two localization positions *y*_−1_ and *y*_+1_. These localization positions share the same location of the molecule along the y-axis. Hence, we can determine the spatial location *y*_0_ in the virtual image by calculating the mean value of *y*_−1_ and *y*_+1_.

We can also perform spectral analysis of individual molecules using all the detected photons. We define the distance between *x*_−1_ in Fig. 1B and *x*_+1_ in Fig. 1C as the spectral shift distance (SSD) (*14*). For individual molecules with longer emission wavelengths (the red plus symbols in Figs. 1B & 1C), their SSD values are larger than the SSDs of molecules with shorter emission wavelengths (the green and blue plus symbols in Figs. 1B & 1C). Therefore, we can distinguish individual molecules based on their distinctive SSDs. To obtain emission spectra of individual molecules, we combine photons from the two spectral images (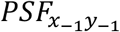 and 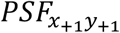) with respect to their spatial locations (*x*_0_, *y*_0_) before spectral fitting, fully utilizing all the collected photons for spectral analysis.

### Single and multi-color SDsSMLM imaging of nanospheres

To test the feasibility of SDsSMLM, we first imaged fluorescent nanospheres (200-nm diameter, F8807, Invitrogen). As a proof of principle, we used a grating (#46070, Edmund Optics) that splits the emitted fluorescence photons into −1^st^, 0^th^, and 1^st^ orders at 22.5%, 28.5%, and 24% transmission efficiency, respectively (Fig. S2). The −1^st^ order and 1^st^ order images are the symmetrically-dispersed spectral images and the 0^th^ order image is the spatial image. Using the 0^th^ order spatial image, we compared the virtual spatial image estimated from the −1^st^ and 1^st^ order spectral images. Figs. 2A & 2B show the two symmetrically-dispersed spectral images and Fig. 2C shows the simultaneously-captured actual spatial images of the nanospheres overlaid with the virtual spatial image. Details of the experiment and the image reconstruction are described in Methods and Materials. We observed that the virtual spatial locations (the green plus symbols in Fig. 2C) of nanospheres estimated from the spectral images agree well with the PSFs and further with the directly obtained spatial locations (the magenta circle symbols). The accuracy of the nanosphere in the highlighted region in Fig. 2C is 4.99 nm. (The average accuracy of five nanospheres is 20.59 nm with standard deviation of 12.71 nm). The magnified views of the highlighted region in Fig. 2C are shown in Figs. 2D-2F. Note that each localization is rendered using a circle with 1-pixel diameter for better illustration in Figs. 2D & 2F. In addition, we numerically corrected the location offset (17.68 ± 23.28 nm and 32 nm ± 9.99 nm (mean ± standard deviation) along the x and y axes respectively) between the virtual and actual spatial locations after image reconstruction.

**Fig. 2.**
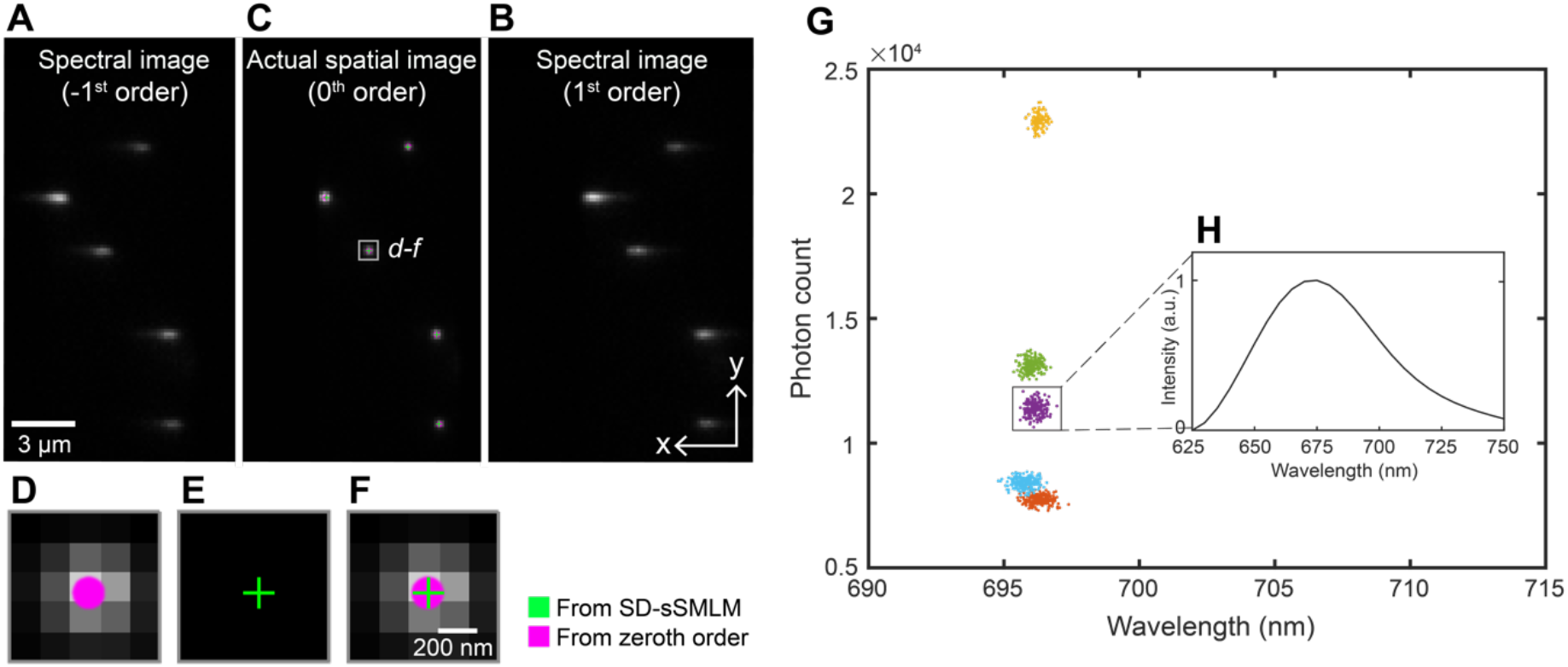
Single-color SDsSMLM imaging of nanospheres. (**A** to **C**) The first frame of the simultaneously captured spectral images and actual spatial image of nanospheres, corresponding to the −1^st^, 1^st^, and 0^th^ orders, respectively; (**D, E**) Magnified views to compare virtual and actual spatial images of the region highlighted by the white box in panel C; **(F)** The corresponding overlaid image of the virtual and actual spatial images; (**G**) The scatter plot of the photon count versus the spectral centroid from five nanospheres; (**H**) The averaged spectrum of one nanosphere from 200 frames, corresponding to the purple cluster in panel G.

We characterized the spectroscopic signatures of the nanospheres using the weighted spectral centroid method (*7*, *23*). Fig. 2G shows the scatter plot of photon count versus spectral centroid for five nanospheres. We observed a narrow spectral centroid distribution of five nanospheres centered at 696 nm with a spectral precision of 0.35 nm. Fig. 2H shows the averaged spectrum of one of the nanospheres from 200 frames (purple cluster in Fig. 2G.)

Besides functional imaging based on spectral analysis (*5*, *10*, *24*), sSMLM allows multi-color imaging with theoretically unlimited multiplexing capability. The multiplexing capability is predominantly determined by the spectral separation of selected dyes and the spectral precision under given experimental conditions (*1*, *7*, *8*). We validated this capability in SDsSMLM using two types of nanospheres (200-nm diameter, F8806 and F8807, Invitrogen). Experimental details are described in Methods and Materials. Figs. 3A & 3B show the first frame of the simultaneously recorded spectral images. While estimating the spatial locations of individual molecules (Fig. 3C), we successfully classified different types of nanospheres based on their spectral centroid distribution (Fig. 3D). The red and blue colors in Fig. 3C correspond to spectral centroids of the crimson nanospheres (centered at 690.6 nm with the spectral precision of 0.48 nm) and for the far-red nanospheres (centered at 696.5 nm with the spectral precision of 0.53 nm), respectively in Fig. 3D.

**Fig. 3.**
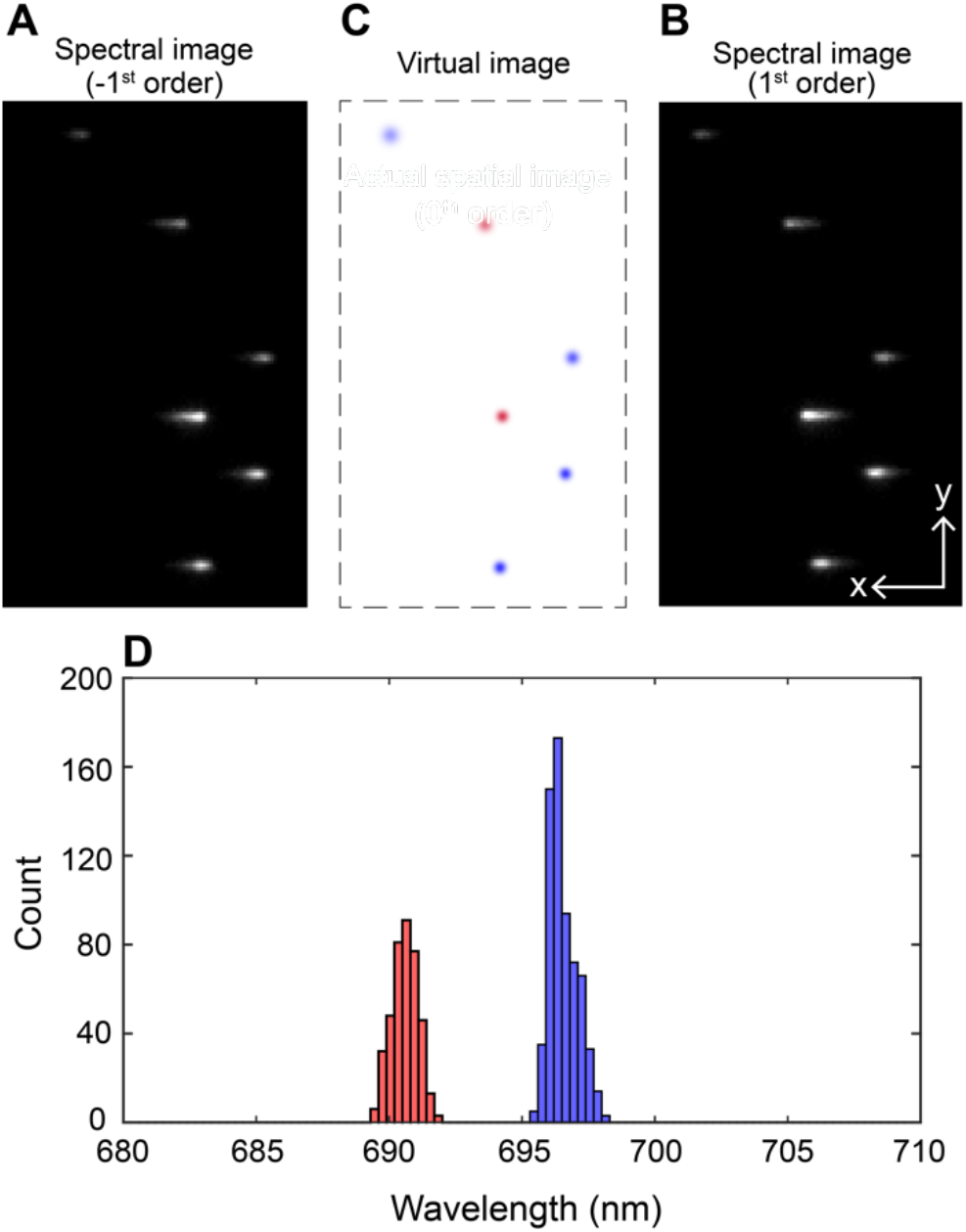
Multi-color SDsSMLM imaging of nanospheres. (**A, B**) The first frame of the simultaneously captured spectral images of nanospheres from the −1^st^ and 1^st^ spectral channels, respectively; (**C**) Calculated virtual spatial image of the nanospheres; (**D**) The spectral centroid distribution of individual nanospheres. The red and blue color in panel C respectively correspond to the spectral centroids of 690.6 nm and 696.5 nm with the spectral precision of 0.48 nm and 0.53 nm.

### Numerical simulation and experimental validation of localization precision in SDsSMLM

In SDsSMLM, collected photons are dispersed into more pixels in spectral image compared with in spatial image (*7*, *23*). Thus, spatial precision of PSF in spectral images (−1^st^ and 1^st^ orders) is more sensitive to noise contribution than spatial precision of PSF in spatial image (0^th^ order). Such spatial precision is not only affected by the number of collected photons and background, but also experimental parameters in the spectral channel, such as spectral dispersion (SD) (*23*) and full width at half-maximum (FWHM) of the emission spectrum. Through numerical simulation, we investigated the influence of SD and FWHM of the emission spectrum on spatial precision as well as spectral precision under different experimental conditions. We further compared the spatial and spectral precisions in SDsSMLM and sSMLM both numerically and experimentally using QDs. The experimental details are described in Methods and the Materials.

We compared achievable spatial and spectral precisions under different SD values (from 1 to 20 nm/pixel) and FWHM values (from 20 to 60 nm), where the total photon count is 1000 in SDsSMLM and sSMLM. In sSMLM, we set the splitting ratio between the spatial (0^th^ order) and spectral (1^st^ order) channels to be 1:3, following previously reported experimental conditions (*3*, *6*–*8*, *11*). We approximated the emission spectrum shape as a Gaussian function. Details of the numerical simulation are described in Methods and Materials. Fig. 4A shows a 2D contour map of the estimated spatial precision of SDsSMLM. Overall, larger SD and narrower FWHM favor high spatial precision. They also favor high spectral precision (Fig. 4B and Fig. S3B). These trends are fundamentally governed by contributions from various noises, such as the signal shot noise, the background shot noise, and the readout noise (*16*, *25*, *26*), and they agree well with analytical solutions, especially for spectral precision (*23*). In comparison, sSMLM shows a uniform spatial precision regardless of SD and FWHM (Fig. S3A). This is because that information in the spectral image in sSMLM is only used for spectral analysis, which is independent from and does not contribute to spatial localization.

**Fig. 4.**
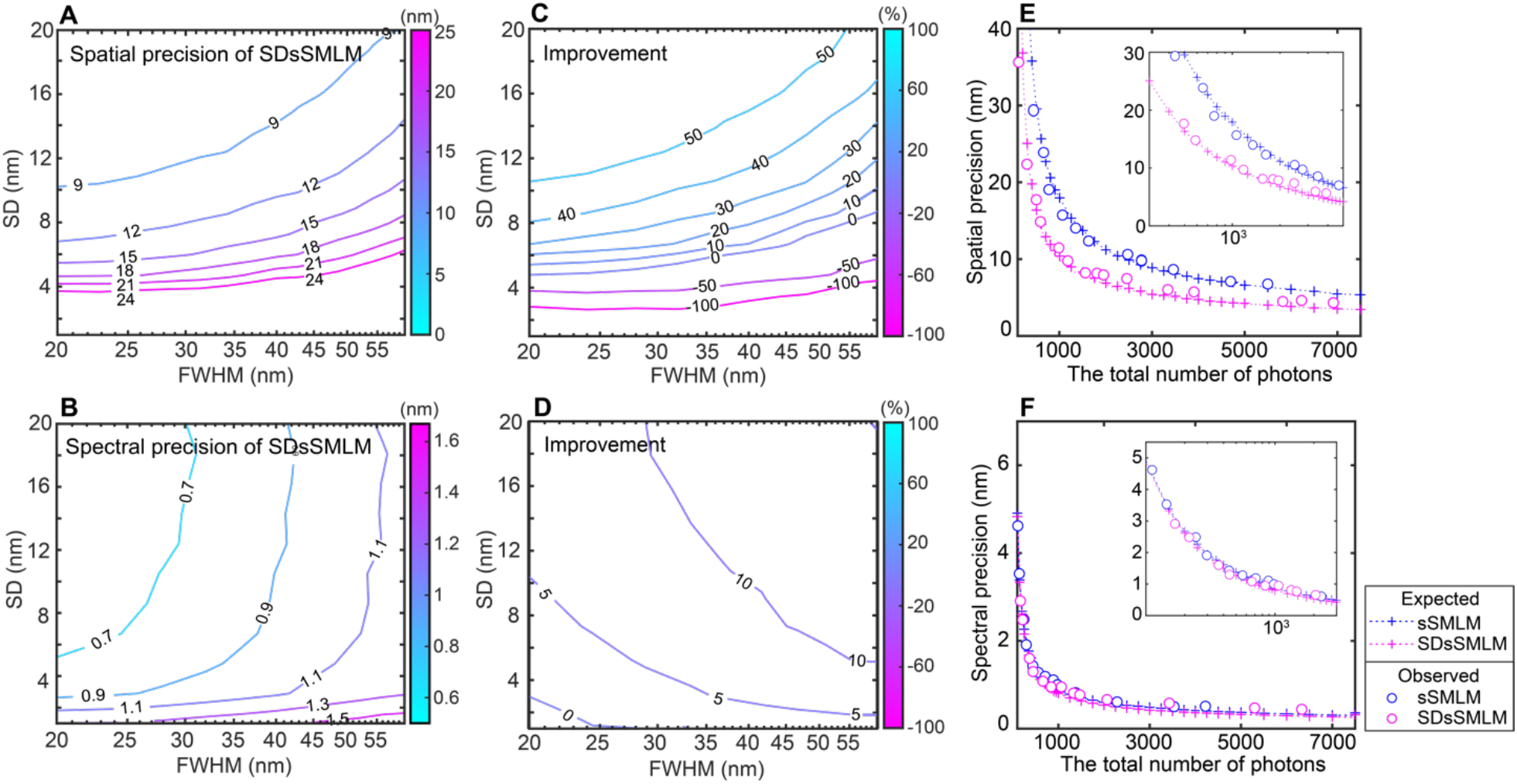
Comparing influences of SD and FWHM of the emission spectrum on the spatial and spectral precisions between SDsSMLM and sSMLM. (**A, B)** Contour map of spatial and spectral precisions under varying SD and FWHM in SDsSMLM; (**C, D**) Contour map of improvements in spatial and spectral precisions in SDsSMLM comparing with sSMLM; (**E,F**) Spatial and spectral precisions of SDsSMLM and sSMLM as a function of the number of photons at 10.5-nm SD and 35-nm FWHM. The magenta and blue colors represent SDsSMLM and sSMLM, respectively. The plus and circle symbols represent the theoretically and experimentally estimated precisions respectively.

Figs. 4C & 4D show improvements in spatial and spectral precisions, respectively, with respect to SD and FWHM in SDsSMLM as compared with sSMLM. For example, at 10.5-nm SD and 35-nm FWHM, which represent the experimental conditions in imaging QDs, SDsSMLM shows approximately 42% (from 17.93 to 10.34 nm) and 10% (from 0.90 to 0.81 nm) higher spatial and spectral precisions, comparing with sSMLM. In particular, SDsSMLM offers a relatively uniform improvement, approximately 10%, in the spectral precision overall. This improvement is proportional to the square root of the ratio of the number of photons allocated to the spectral channel between SDsSMLM and sSMLM (Figs. 4D). We further estimated the achievable spatial and spectral precisions when the number of photons increases. As shown in Figs. 4E & 4F, the theoretical estimations are in good agreement with experimental results using QDs. In addition, we investigated the influence of splitting ratios on the spatial and spectral precisions (Fig. S4).

### Multi-color SDsSMLM imaging of COS7 cells

We demonstrated multi-color imaging capability of SDsSMLM using fixed COS7 cells. We selected Alexa Fluor 647 (AF647) and CF680, which emit at wavelengths only ~30 nm apart (Fig. 5A), to label mitochondria and peroxisomes, respectively (*1*, *8*). To classify them, we used different spectral bands based on the spectral centroid distribution (*7*, *8*): the first band from 682 nm to 694 nm for AF647 and the second band from 699 nm to 711 nm for CF680, as respectively highlighted by the yellow and cyan colors in Fig. 5B. We visualized the colocalization of mitochondria (yellow) and peroxisomes (cyan) (Fig. 5C). We also imaged microtubules labeled with AF647 (magenta) and mitochondria labeled with CF680 (green) (Fig. 5D). By measuring the FWHM of a segment of imaged microtube (dashed square in Fig. 5D), we estimate the spatial resolution of SDsSMLM to be 66 nm as shown in Fig. 5E. Also, we observed that the minimum resolvable distance between two tubulin filaments is within the range of 81-92 nm from multiple Gaussian fittings of the intensity profiles (Figs 5F & 5G). Using a Fourier ring correlation method (FRC) method (*27*, *28*), we also evaluated the resolution of another reconstructed image (Fig. 5C) that visualizes mitochondria and peroxisomes. The FRC curve estimated a resolution of 111 nm (Fig. S5) at a threshold level of 1/7.

**Fig. 5.**
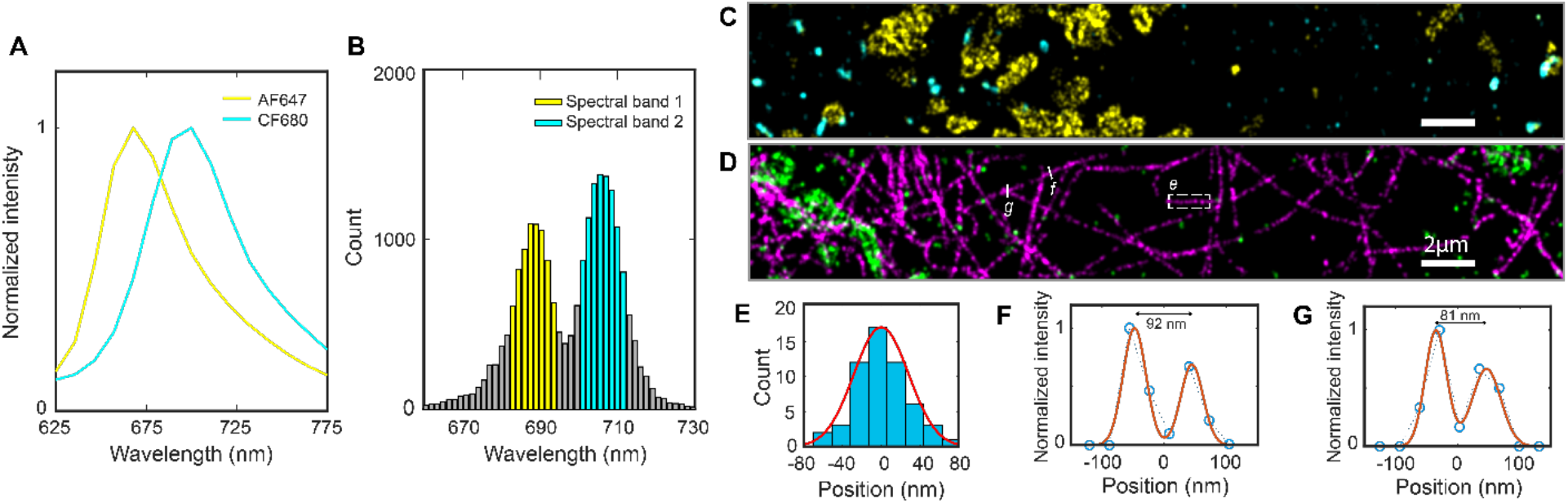
Multi-color SDsSMLM imaging of fixed COS7 cells. (**A**) Normalized emission spectra of AF647 (yellow) and CF680 (cyan); (**B**) Spectral centroid distributions of AF647 and CF680. The first spectral band (yellow) covers 682 nm to 694 nm for AF647 and the second spectral band (cyan) covers 699 nm to 711 nm for CF680; (**C**) Reconstructed multi-color SDsSMLM image of mitochondria (yellow) and peroxisomes (cyan); (**D**) Reconstructed multi-color SDsSMLM image of microtubules labeled with AF647 (magenta) and mitochondria labeled with CF680 (green); (**E**) Histogram of the cross-section highlighted by the white-dashed box in panel D. (**F, G**) Intensity profiles of two imaged tubulin filaments highlighted by the white-solid lines in panel D.

In addition, we quantified the utilization ratio, which is defined as the ratio between the number of localizations allocated into each spectral band to the total number of localizations, in the reconstructed image (Fig. 5C). We calculated the utilization ratio in SDsSMLM using both spectral images. We also calculated the utilization ratio by using only one spectral image (1^st^ order channel), which mimics conventional sSMLM with a 1:1 splitting ratio between the spatial and spectral channels, for comparison. We obtained a 17.4% improvement in utilization ratio in SDsSMLM, on average for the two spectral channels, comparing with sSMLM (Fig. S6). This result demonstrates that SDsSMLM benefits from improved spectral precision by fully utilizing all collected photons for spectral analysis, which, subsequently, leads to improved spectral classification for multi-color imaging.

### 3D single particle tracking

We added 3D imaging capability to SDsSMLM through biplane imaging, similar to what we reported in sSMLM (*7*). Since SDsSMLM already has two symmetrically-dispersed spectral channels, we can efficiently implement biplane imaging by introducing an extra optical path length in one channel. As shown in Fig. 6A, we added a pair of mirrors in the 1^st^ order spectral channel in front of the EMCCD camera to generate such optical path length difference. This optical path length difference introduced a 500-nm axial separation between the imaging planes of the two spectral channels. As a result, individual molecules are imaged with differently PSF sizes in the two spectral images according to their axial locations. By measuring the ratio between the sizes of the PSFs captured in the two spectral images, we can determine the axial coordinate of each molecule through an axial calibration curve. The full description of biplane SDsSMLM image reconstruction is described in Methods and Materials.

**Fig. 6.**
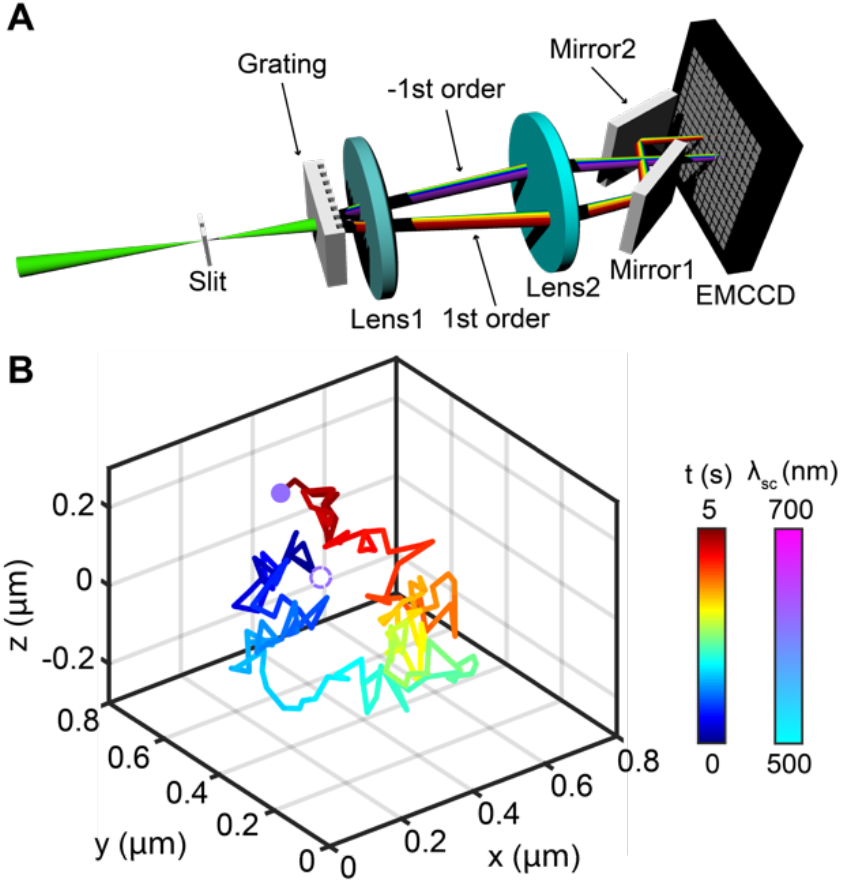
3D biplane SDsSMLM. (**A**) Schematic of the experimental system; (**B**) Imaged 3D trajectory of a single QD color-coded with respect to acquisition time (solid line). The QDs at the first and last frames are highlighted by color-coded circles with respect to measured spectral centroids.

We demonstrated 3D biplane SDsSMLM by tracking individual QDs (777951, Sigma-Aldrich) in a suspension. We tracked the movement of QDs for 5 s. We recorded 160 frames with an exposure time of 5 ms at a frame rate of 30 Hz. Fig. 6B shows the 3D trajectory of one QD, color-coded along the time (represented by the line). The QD locations at the first and last frames are highlighted by the color-coded circles according to the measured spectral centroids. We observed that the spectral centroids remained near 614 nm throughout tracking period with a spectral precision of 1.5 nm (Fig. S7A). We approximated the diffusion coefficient from the 3D trajectory using *D* = *MSD* / 6*t*, where *MSD* is the mean squared displacement and *t* is frame acquisition time (*29*). The calculated diffusion coefficient is 0.021 μm^2^/s. These results demonstrate the capability of 3D biplane SDsSMLM to precisely reconstruct the 3D spatial and spectral information of single molecules in SPT.

## Discussion

We demonstrated that SDsSMLM acquires both spatial and spectral information of single molecules from two symmetrically-dispersed spectral images without capturing the spatial image. SDsSMLM maintains the highest achievable spectral precision per each emitter in a given experimental condition as it fully uses all collected photons for spectral analysis. In addition, it addresses an inherent trade-off between spatial and spectral precisions by sharing all collected photons in both spatial and spectral channels. We observed that SDsSMLM achieved 10.34-nm spatial and 0.81-nm spectral precisions with 1000 photons, which correspond to 42%, approximately doubled photon enhancement, and 10% improvements in the spatial and spectral precisions, respectively, comparing with sSMLM using a 1:3 ratio between the spatial and spectral channels.

We applied SDsSMLM to multi-color imaging and 3D SPT. It should be noted that these experimental demonstrations were based on a grating that splits the beam to the −1^st^ and 1^st^ orders with the efficiency of 22.5% and 24%, respectively. Thus, only approximately half of photons of the emitted fluorescence were used for image reconstruction in multi-color imaging. Consequently, current implementation of SDsSMLM had a reduced image resolution. This can be improved by replacing this grating with a new phase grating, which can significantly suppress the 0^th^ order and maximize the transmission efficiency only at the −1^st^ and 1^st^ orders, with a relatively high total transmission efficiency expected to be more than 85%. In addition, the resolution can be further improved by using a larger SD and narrower FWHM as SDsSMLM favors a large SD and narrow FWHM for higher spatial precision. However, an extremely low SD may compromise one of the benefits of SDsSMLM for functional studies that involve resolving minute spectroscopic features in single-molecule spectroscopy. This suggests that SDsSMLM requires a careful dye selection and system optimization to achieve desired spatial and spectral precisions.

The field-of-view (FOV) in SMLM is mainly determined by the objective lens, the field of illumination, and the active area of the camera. For sSMLM equipped with a grating-based spectrometer, the FOV is further restricted by the diffraction angle of the 1^st^ order of the grating, which determines the separation between the spatial and spectral images. In this work, our FOV was restricted to ~30×5 μm^2^ as we also captured the 0^th^ order to compare the virtual and actual spatial images. This constraint can be relaxed in the future by a customized grating, which suppresses the 0^th^ order. In this case, the FOV primarily depends on the separation between the −1^st^ and 1^st^ orders, which could increase the FOV by at least two-fold. Also, it can be further addressed in 3D biplane SDsSMLM by manipulating two diffraction orders separately.

In 3D biplane SDsSMLM, the PSFs of individual molecules in the spectral images are blurred when they are at out of focus planes. This does not allow for a detailed spectral analysis. However, the spectral centroid calculated by the intensity-weighted method still can be used to separate two dyes with slightly different fluorescence spectra (*7*). In addition, we have small differences in the magnification and the SD between the two spectral images caused by their different path lengths. However, the spectral centroid is not significantly affected by these issues and is sufficient for extracting spectroscopic signatures of individual molecules. We numerically corrected them before image reconstruction and spectral analysis. And, we observed the spectral precision of 1.5 nm throughout tracking period in a given experimental conditions: The signal level is ~6700 photons and the background level is ~800 photons in total.

## Materials and Methods

### Optical setup and image acquisition for SDsSMLM imaging

We performed all experiments using a home-built SDsSMLM system, which is based on an inverted microscope body (Eclipse Ti-U, Nikon) (Fig. S1). We used a 640-nm laser to excite nanospheres, AF647, and CF680 and used a 532-nm laser (Excelsior One-532-200, Spectra Physics) to excite QDs. The laser beam was reflected by a dichroic filter (FF538-FDI01/FF649-DI01-25X36, Semrock) and focused onto the back aperture of an oil immersion objective lens (CFI Apochromat 100X, NA=1.49, Nikon). We used a high oblique angle to illuminate the samples. The emitted fluorescence light was collected by the objective lens and focused by the tube lens onto the intermediate image plane after passing through a long pass filter (LPF) (BLP01-532R/647R-25, Semrock). We inserted a slit at the intermediate image plane to confine the FOV and subsequently placed a transmission grating (46070, Edmund Optics) to disperse the emitted fluorescence into the −1^st^, 0^th^, and 1^st^ orders. Then, the dispersed fluorescence emissions were captured by an EMCCD camera (iXon 897, Andor) with a back-projected pixel size of 160 nm after passing through the relay optics (the focal length, *f* = 150 mm, AC508-150-B-ML, Thorlabs).

For SDsSMLM imaging of nanospheres, we acquired 200 frames at a power density of ~0.02 kW/m^2^ with an exposure time of 20 ms. For the experimental validation of localization precision using QDs, we acquired 200 frames while varying signal intensity (photon count) by adjusting EMCCD exposure time and controlling illumination power using a neutral density filter (NDC-50C-4M, Thorlabs). For multi-color SDsSMLM imaging of fixed COS7 cells, we acquired 20000 frames at ~10 kW/cm^2^ with an exposure time of 20 ms. For SPT in 3D, we acquired 160 frames at ~0.02 kW/cm^2^ with an exposure time of 5 ms.

### Image reconstruction for SDsSMLM imaging

For image reconstruction, we first localized two spectral images (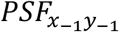 and 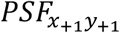) of individual molecules with 2D Gaussian fitting using ThunderSTORM (*21*). Then, using customized MATLAB codes, we classified them into two groups corresponding to the −1^st^ and 1^st^ orders and estimated the spatial locations of pairs of localizations by calculating their mean values. Next, we formed the virtual image (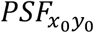) using the estimated spatial locations.

For spectral calibration, we first captured a calibration image using a narrow slit and a fluorescence calibration light source. This calibration image includes multiple spectral lines of the calibration light source in the two spectral images. By integrating the two spectral images along the y-axis, we obtained the multiple emission peaks centered at 487.7, 546.5, and 611.6 nm (Fig. S8A). Then, we obtained a calibration curve by fitting the wavelengths of the emission peaks with their corresponding pixel distances using a linear polynomial function (Fig. S8B). Using the obtained calibration curve, we calibrated the emission spectra of individual molecule pairs. Lastly, we obtained the final emission spectra of individual molecules by combining the two symmetrical emission spectra. Note that the grating provides an approximate linear relationship within the spectral range used in the experiment. In addition, we used three main emission peaks of 487.7, 546.5, and 611.6 nm of a fluorescent calibration lamp for the first experiment demonstration using nanospheres while using two main emission peaks at 620.23 and 603.24 nm of the Neon lamp (6032, Newport) for the rest of the experimental demonstrations.

To characterize spectroscopic signatures of individual molecules, we used spectral centroid, which is calculated by the weighted centroid method (*23*). For all the experimental demonstrations, we estimated the spectral centroid in the same manner except multi-color imaging using nanospheres. Unfortunately, in this experiment, we rarely distinguished two different types of nanospheres based on spectral centroid values as the emission of one of nanospheres (crimson) was partially rejected by the LPF. Thus, we fitted the emission spectrum using Gaussian function and used the emission peak of the fitted spectrum as an approximation of spectral centroid.

For imaging nanospheres, QDs, and fixed cells, we used the spectral window of 650-750 mm, 565-665 nm, and 625-775 nm, respectively. Besides, we rejected blinkings below 500 photons during the spectral analysis in multi-color imaging of fixed cells. In addition, we used an SD of 8.8 nm/pixel in nanosphere imaging and an SD of 10.5 nm in all other experiments.

### Numerical simulation

To simulate two symmetrical spectral images in SDsSMLM, we first generated a spatial image. The spatial image was modeled as a 2D Gaussian function with a sigma value of 0.94 pixel, which represents the experimental conditions: back-projected pixel size of 160 nm and PSF FWHM of 350 nm. Then, we convolved the generated spatial image with the emission spectrum of the dye molecule being simulated (with different FWHM and SD values) to generate a spectral image (1^st^ order). Next, we generated an identical spectral image (−1^st^ order). We modeled various noise sources, such as the signal shot noise, the background shot noise, and the readout noise. The shot and readout noises follow Poisson and Gaussian distributions, respectively. Finally, we generated noise-added spectral images at different signal and noise levels (*23*). We used the readout noise of 1e- and 3000 iterations in all simulations.

We estimated the spatial precision of SDsSMLM using the simulated spectral images. The spatial precision was calculated using the standard deviation of the distribution of the estimated (*x*_0_, *y*_0_) in the virtual spatial image (Fig. 1D). By averaging the two spatial precisions along the x and y axes, we calculated a final spatial precision.(*22*) In addition, we estimated the spectral precision of SDsSMLM using the simulated spectral images. The spectral precision was calculated using the standard deviation from the distribution of the spectral centroid *λ*_*SC*_.

To compare the performance of SDsSMLM with that of sSMLM, we also estimated the spatial precision in sSMLM. For sSMLM, we generated noise-added spatial images at different signal and noise levels. Then, we estimated the spatial precision using standard deviation from the spatial location distributions. Additionally, we estimated the spectral precision of sSMLM. This procedure was essentially the same as described for SDsSMLM, except that only one spectral image corresponding to the 1^st^ order was used to obtain the emission spectrum.

Finally, we compared the spatial and spectral precisions of SDsSMLM with those obtained from sSMLM given varying splitting ratios between the 0^th^ and 1^st^ orders. For fair comparisons, we assumed that SDsSMLM and sSMLM share the same total number of photons. For SDsSMLM, the total photons were split equally between the −1^st^ and 1^st^ orders while sSMLM varied splitting ratios between the 0^th^ and 1^st^ orders.

### Image reconstruction for 3D biplane SDsSMLM

We reconstructed the 3D image in the similar manner as previously described in 3D biplane sSMLM (*7*), except that we used one symmetrically-dispersed spectral image (−1^st^ order) instead of the spatial image (0^th^ order), together with another spectral image (1^st^ order) for biplane imaging. We first captured a 3D calibration image using QDs. This image contains a few samples in both spectral images at different depths. The QDs were scanned from −1.5 μm to +1.5 μm along the z axis with a step size of 25 nm. Next, we obtained one dimensional (1D) PSF_y_s by integrating the spectral images along the x axis. Then, we measured the FWHM of two 1D PSF_y_s and estimated their ratio (Fig. S7B). We used this ratio to calibrate the axial coordinate of each molecule.

For all detected molecules, we calculated the PSF_y_s FWHM ratio of the two symmetrically-dispersed spectral images and estimated their axial coordinates using the aforementioned 3D calibration curve. Then, we combined them with the 2D spatial and spectral information to generate a 4D array for individual molecules.

### Sample preparation for SDsSMLM imaging

We prepared nanosphere samples for single and multi-color SDsSMLM imaging according to the following steps. No. 1 borosilicate-bottom 8-well Lab-Tek Chambered cover glasses were rinsed with PBS, coated with poly-l-lysine (PLL, P8920, Sigma-Aldrich) for 1 hour, and washed with PBS 3 times. Nanospheres (200 nm diameter; F8806 and F8807, Invitrogen) were diluted 10^4^ times with a cross-linking buffer containing EDC (1 mg mL^−1^, 1-ethyl-3-(3-dimethylaminopropyl) carbodiimide hydrochroride) and NHS (1 mg mL^−1^, N-Hydroxysuccinimide) in 50 mM MES buffer (2-(N-morpholino) ethanesulfonic acid, pH = ~6, 28390, ThermoFisher). 200 μL of the cross-linking buffer with nanospheres was added to the PLL-coated cover glass for 5 min for surface immobilization. The cover glass was rinsed with PBS and dried under filtered air. Then, a drop of antifade mounting medium (P36965, Invitrogen) was added to a cover slip. The cover glass with samples were mounted on the cover slip by sandwiching the samples between them and then sealed with the black nail polish.

We prepared QD sample according to the following steps. QDs (777951, Sigma-Aldrich) were diluted 10^4^ times in water. 400 μL of the QD solution with a concentration of 0.5 μg mL^−1^ was deposited onto the cover glass using Laurell WS-650SZ-23NPPB spin-coater at 2000 rpm for 1 min. The cover glass with samples was mounted on the cover slip by sandwiching the samples between them and then sealed with the black nail polish.

We prepared fixed COS7 cell samples for multi-color SDsSMLM imaging according to the following steps. COS7 cells (ATCC) were maintained in Dulbecco’s Modified Eagle Medium (DMEM, Gibco/Life Technologies) supplemented with 2 mM L-glutamine (Gibco/Life Technologies), 10% fetal bovine serum (Gibco/Life Technologies), and 1% penicillin and streptomycin (100 U mL^−1^, Gibco/Life Technologies) at 37°C with 5% CO2. Cells were plated on a cover glass with about 30% confluency. After 48 hours, we rinsed cells with phosphate buffered saline (PBS), then fixed the cells with 3% paraformaldehyde and 0.1% glutaraldehyde in PBS for 10 min at room temperature. After washing with PBS twice, the cells were quenched with 0.1% sodium borohydride in PBS for 7 min and rinsed twice with PBS. The fixed cells were permeabilized with a blocking buffer (3% bovine serum albumin (BSA), 0.5% Triton X-100 in PBS for 20 min), followed by incubation with the primary antibodies in the blocking buffer for 1 hour at room temperature. For multi-color imaging of mitochondria and peroxisomes, the primary antibodies used in the study are mouse anti-TOM20 directly labeled with AF647 (2.5 μg mL^−1^, sc-17764-AF647, Santa Cruz) and rabbit anti-PMP70 (1:500 dilution, PA1-650, ThermoFisher). The samples were washed three times with washing buffer (0.2% BSA, 0.1% Triton X-100 in PBS) for 5 min and incubated with the secondary antibodies labeled with CF680 (2.5 μg mL^−1^ donkey anti-rabbit IgG -CF680) for 40 min. For multi-color imaging of microtubules and mitochondria, the primary antibodies used in the study were sheep anti-tubulin (2.5 μg mL^−1^, ATN02, Cytoskeleton) and mouse anti-TOM20 (2.5 μg mL^−1^, sc-17764, Santa Cruz). After washing with washing buffer three times for 5 min and incubated with the secondary antibodies labeled with AF647 and CF680 (2.5 μg mL^−1^ donkey anti-sheep IgG-AF647, anti-mouse IgG-CF680) for 40 min. The dyes were conjugated to the IgG following a literature protocol (degree of label = ~1) (*30*). The cells were then washed with PBS three times for 5 min and stored at 4°C. An imaging buffer (pH = ~8.0, 50 mM Tris, 10 mM NaCl, 0.5mg mL^−1^ glucose oxidase (G2133, Sigma-Aldrich), 2000 U/mL catalase (C30, Sigma-Aldrich), 10% (w/v) D-glucose, and 100 mM cysteamine replaced PBS) was replacing PBS before image acquisition.

Sample preparation for 3D SPT using a free-diffusing QD are described as the following. A QD solution of 0.5 μg mL^−1^ in water was mix with Glycerol (v/v = 1:9) and vortexed for 10 s. Then 50 μL of the final solution was added onto a cover glass immediately. The free-diffusing single QD was then observed and tracked.

## Acknowledgments

This work was supported in part by NSF grants CBET-1706642 and EFMA-1830969; NIH grants R01EY026078 and R01EY029121. Ki-Hee Song is supported by the Christine Enroth-Cugell and David Cugell Graduate Fellowship in Biomedical Engineering and Visual Neuroscience.

