## Supplementary figures for "Symmetrically-dispersed spectroscopic single-molecule localization microscopy"

This PDF file includes:

**Fig. S1**. Schematic of SDsSMLM system.
**Fig. S2**. Schematic of a grating used for the experimental demonstrations.
**Fig. S3**. Influences of SD and FWHM of the emission spectrum on the spatial and spectral precisions in sSMLM.
**Fig. S4**. Additional comparisons of the spatial and spectral precisions between SDsSMLM and sSMLM.
**Fig. S5**. FRC curve of the reconstructed multi-color image.
**Fig. S6**. Analysis of the utilization ratio of the reconstructed multi-color image.
**Fig. S7**. 3D SPT.
**Fig. S8**. Spectral calibration information.

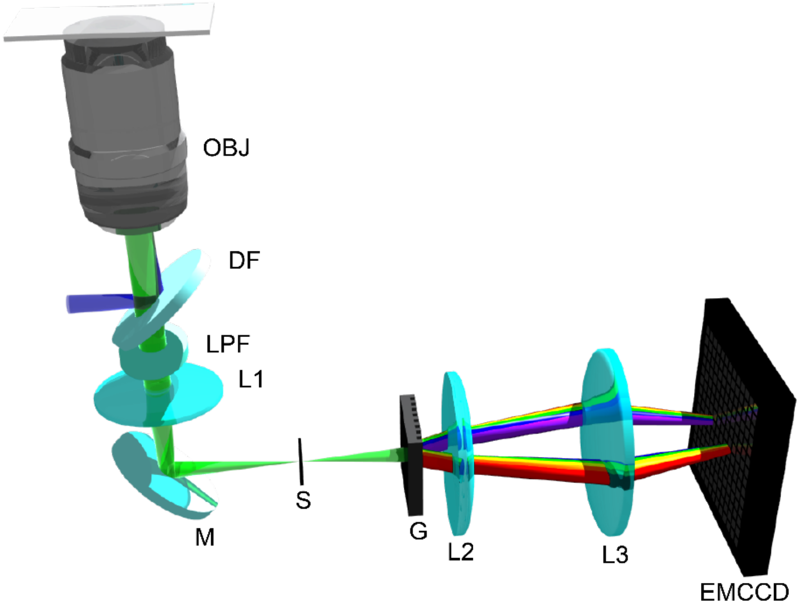


**Fig. S1**. Schematic of SDsSMLM system. OBJ: Objective Lens; DF: Dichroic Filter; LPF: Long Pass Filter; L: Lens; M: Mirror; S: Slit; G: Grating; EMCCD: Electron Multiplying Charge Coupled Device.

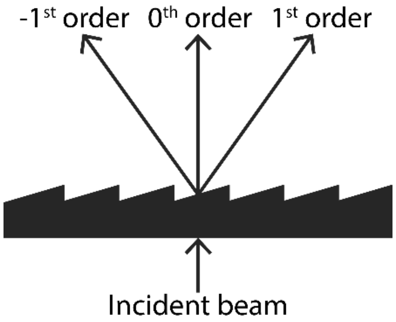


**Fig. S2**. Schematic of a grating used for the experimental demonstrations.

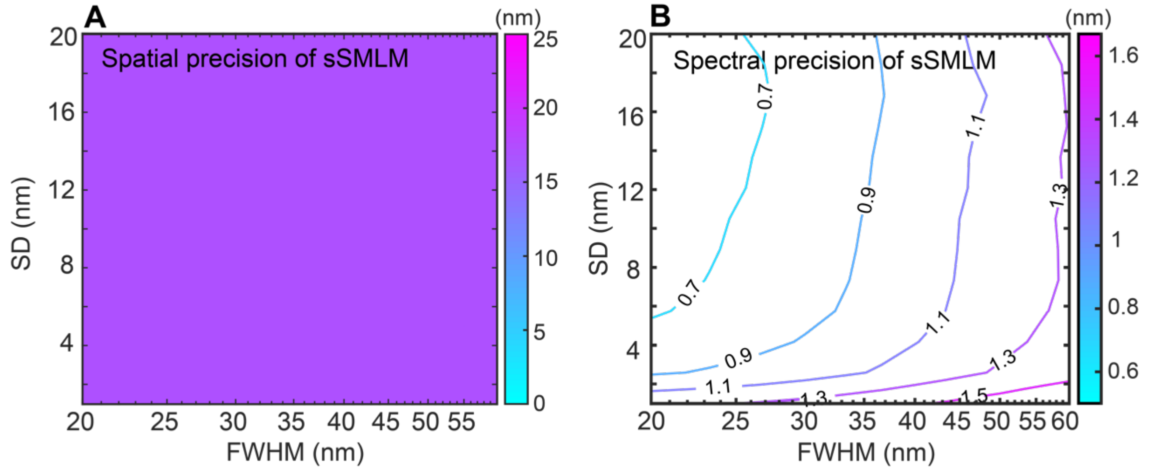


**Fig. S3**. Influences of SD and FWHM of the emission spectrum on the spatial and spectral precisions in sSMLM. (**A, B**) Contour map of spatial and spectral precisions under varying SD and FWHM.

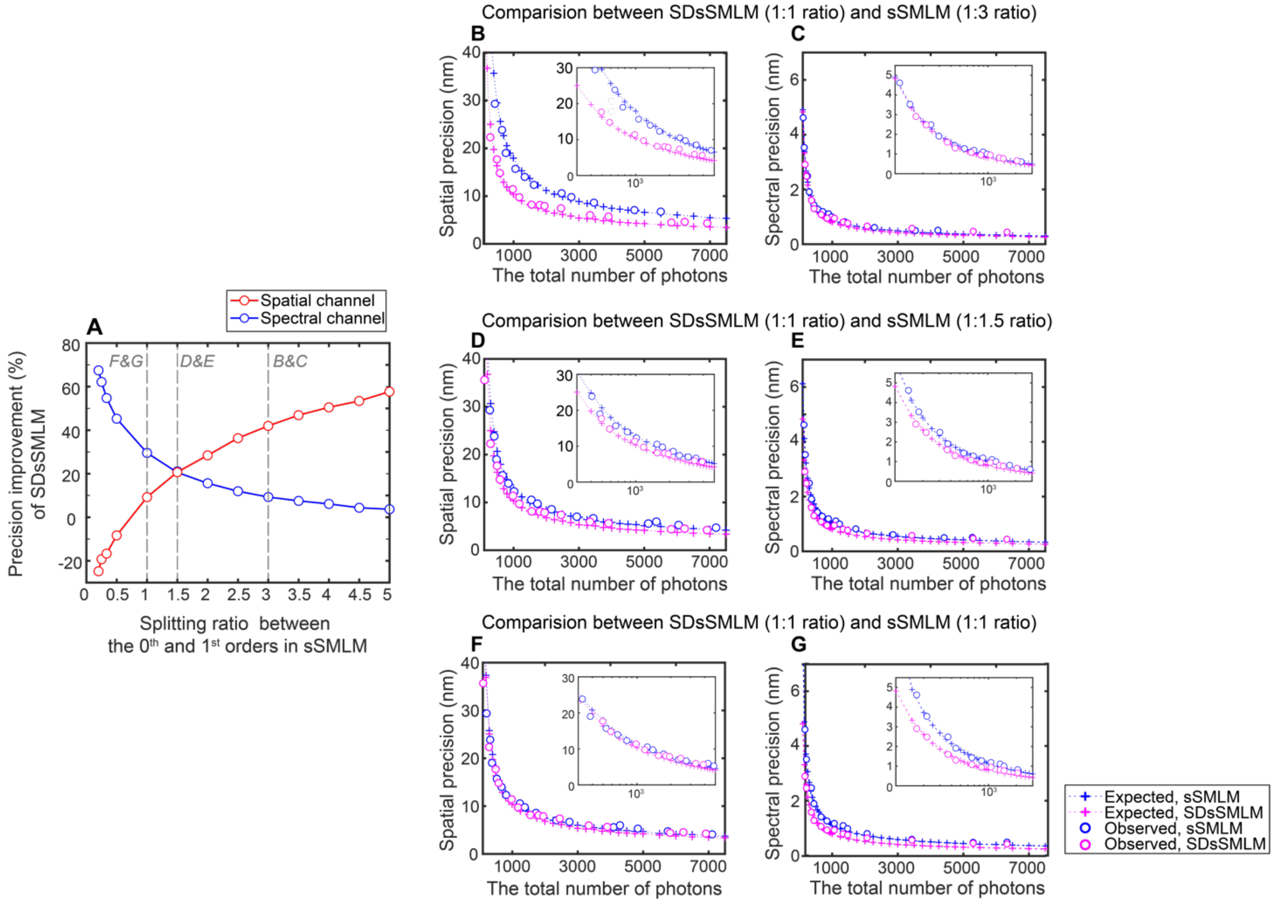


**Fig. S4**. Additional comparisons of spatial and spectral precisions between SDsSMLM and sSMLM. (**A**) Expected precision improvement of SDsSMLM when splitting ratio between the 0$^{th}$ and 1$^{st}$ orders in sSMLM varies. The signal level is 1000 photons. At the splitting ratio of (**B**, **C**) 1:3, (**D**, **E**) 1:1.5, and (**F, G**) 1:1 for sSMLM, the achievable spatial and spectral precisions as a function of the number of photons. SDsSMLM achieves 42% (from 17.93 nm to 10.34 nm) and 10% (from 0.90 nm to 0.81 nm) improvements in spatial and spectral precisions, respectively, compared with sSMLM featuring a 1:3 ratio; (1) 19% spatial (from 12.73 nm to 10.34 nm) and 21% spectral (from 1.03 nm to 0.81 nm) precision improvements compared to sSMLM with a 1:1.5 ratio; (2) 10% spatial (from 11.42 nm to 10.34 nm) and 30% spectral (from 1.15 nm to 0.81 nm) precision improvements compared to sSMLM with a 1:1 ratio.

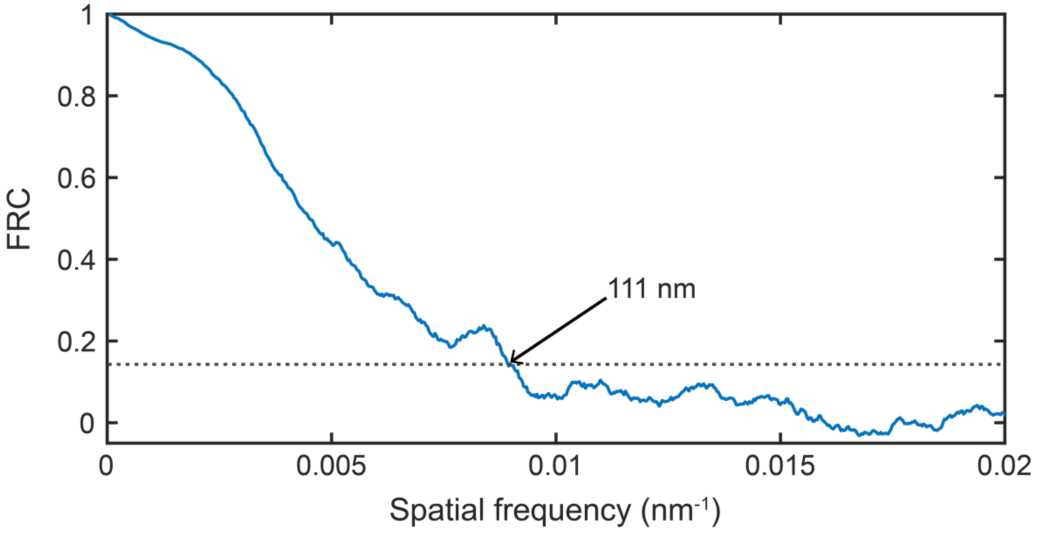

**Fig. S5**. FRC curve of the reconstructed multi-color image shown in Fig. 5C.

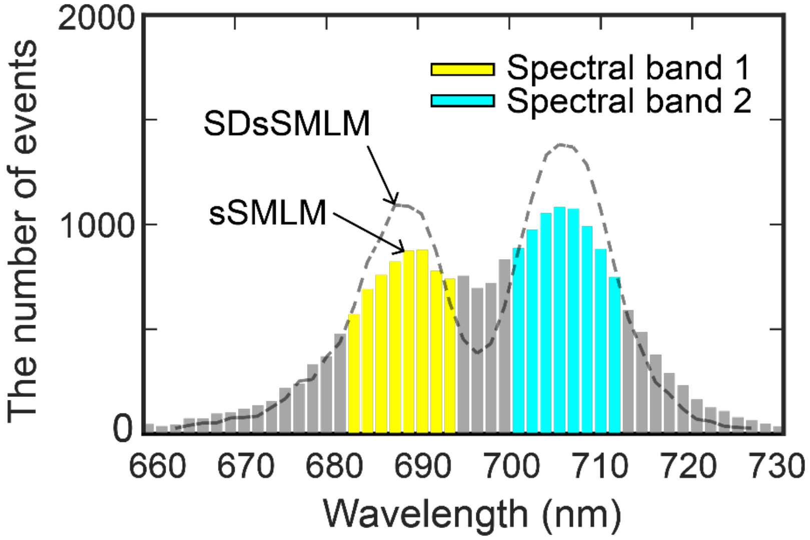


**Fig. S6**. Analysis of utilization ratio for the reconstructed multi-color image shown in Fig. 5C. The histogram represents the spectral centroid distribution estimated from only one spectral image corresponding to the 1$^{st}$ order. This case reasonably mimics conventional sSMLM with a 1:1 splitting ratio between the spatial and spectral channels. The dashed line shows the profile of the spectral centroid distribution estimated from two spectral images in the SDsSMLM case. The number of localizations allocated to each spectral band was increased from 6074 to 7119 for the first spectral channel and from 7759 to 9124 for the second spectral channel, which correspond to 17.2% and 17.6% improvements in the utilization ratio, respectively.

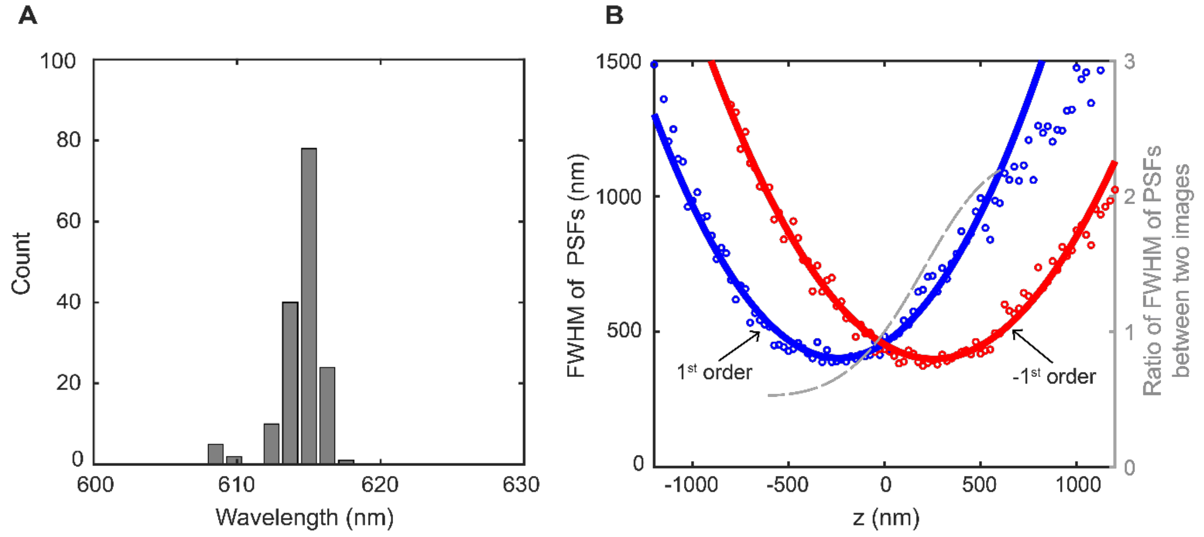


**Fig. S7**. (**A**) Histogram of spectral centroids of the QD during tracking; (**B**) 3D calibration curve. The blue and red solid lines indicate the FWHM of PSFs in the $1^{st}$ and $-1^{st}$ orders respectively. The gray color dash line represents the corresponding ratio.

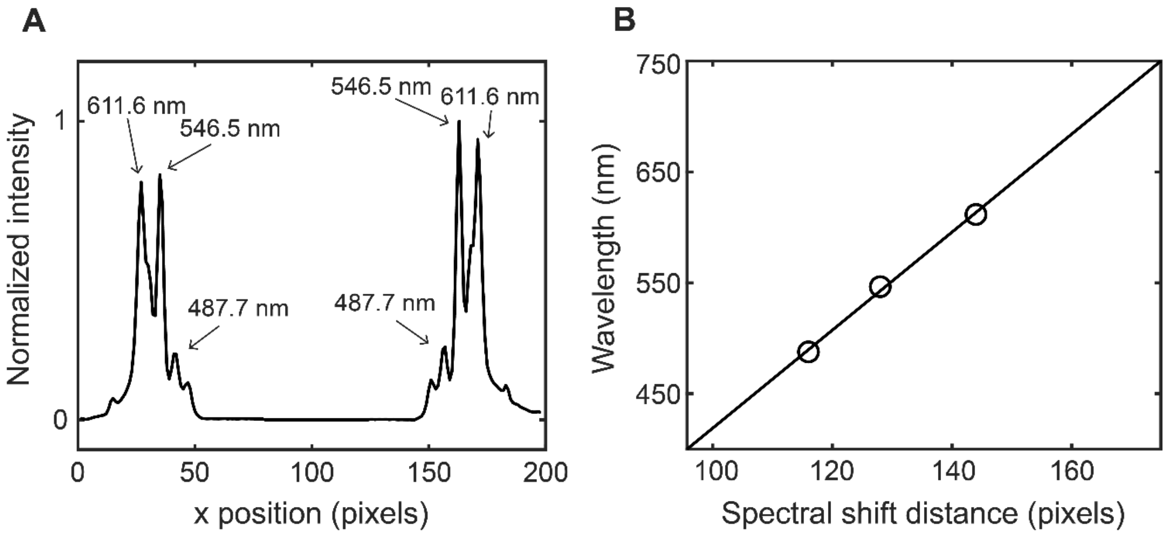


**Fig. S8**. Spectral calibration information. (**A**) Emission peaks of the calibration source centered at 487.7 nm, 546.5 nm, and 611.6 nm; (**B**) Calibration curve obtained by fitting the wavelengths with their corresponding pixel distances using a linear polynomial function.
